# Cerebellar Dentate Connectivity Across Adulthood: A Large-Scale Resting State Functional Connectivity Investigation

**DOI:** 10.1101/2021.02.24.432761

**Authors:** Jessica A. Bernard, Hannah K. Ballard, T. Bryan Jackson

**Author notes:** Corresponding Author Jessica A. Bernard, PhD, Department of Psychological and Brain Sciences, Texas A&M University.

## Abstract

Cerebellar contributions to behavior in advanced age are of interest and importance, given its role in motor and cognitive performance. There are differences and declines in cerebellar structure in advanced age and cerebellar resting state connectivity is lower. However, the work on this area to date has focused on the cerebellar cortex. The deep cerebellar nuclei provide the primary cerebellar inputs and outputs to the cortex, as well as the spinal and vestibular systems. Dentate networks can be dissociated such that the dorsal region is associated with the motor cortex, while the ventral aspect is associated with the prefrontal cortex. However, whether dentato-thalamo-cortical networks differ across adulthood remains unknown. Here, using a large adult sample (n=590) from the Cambridge Center for Ageing and Neuroscience, we investigated dentate connectivity across adulthood. We replicated past work showing dissociable resting state networks in the dorsal and ventral aspects of the dentate. In both seeds, we demonstrated that connectivity is lower with advanced age, indicating that connectivity differences extend beyond the cerebellar cortex. Finally, we demonstrated sex differences in dentate connectivity. This expands our understanding of cerebellar circuitry in advanced age and underscores the potential importance of this structure in age-related performance differences.

## Introduction

In the last century we have seen a rapid increase in the human lifespan and demographic shifts, wherein aging individuals make up an increasingly large proportion of the population. As such, understanding the aging brain and differences in behavior is of increasing importance for the quality of life and health of older adults (OA). Even in the best cases of healthy aging without known neurological or somatic pathology, differences in both cognitive and motor performance are observed in OA (Park and Reuter-Lorenz 2009; Seidler et al. 2010a; Reuter-Lorenz and Park 2014; Cabeza et al. 2018). Understanding normative differences in brain function and organization in young adults (YA) and OA provides critical insights into the underpinnings of the behavioral changes observed in healthy aging. Furthermore, understanding normative aging and the associated brain and behavioral trajectories across adulthood stand to provide important insights and points of comparison for investigations of age-related diseases, such as Alzheimer’s Disease.

With this in mind, there have been significant advances in our understanding of age differences in brain structure and function, as well as changes over time. We know now that even in healthy aging there are marked differences in brain structure (e.g., Raz et al. 1997, 1998, 2013; Bernard and Seidler 2013; Bernard et al. 2015), patterns of functional activation (Reuter-Lorenz et al. 1999; Cabeza 2002; Cabeza et al. 2002, 2018; Reuter-Lorenz and Cappell 2008), and network connectivity (Andrews-Hanna et al. 2007; Tomasi and Volkow 2012; Ferreira and Busatto 2013; Bo et al. 2014; Ferreira et al. 2016). However, to this point, much of the research on understanding the aging brain and behavior has focused on cortical contributions, and the cerebellum has been relatively understudied. The human cerebellum is a neuronally dense structure with extensive folding, such that its surface area is 78% of that of the cerebral cortex (Sereno et al. 2020), and it contributes to both motor and non-motor behaviors (Schmahmann and Sherman 1998; Stoodley et al. 2012; Schmahmann 2018; King, Hernandez-Castillo, Sereno, et al. 2019). It is connected to the cortex through closed-loop circuits with the thalamus, that have been mapped both in non-human and human primates (Kelly and Strick 2003; Ramnani 2006; Strick et al. 2009; Salmi et al. 2010; Bernard et al. 2016). Further, at rest, when investigating cerebello-cortical functional connectivity there are distinct connections with the cortex when mapped using individual cerebellar lobules or distinct proscribed subregions (Krienen and Buckner 2009; O’Reilly et al. 2010; Bernard et al. 2012). The cerebellum can also be parcellated in accordance with known cortical resting-state networks (Buckner et al. 2011). More recently, it has been suggested that the organization of the cerebellum follows functional gradients based on connectivity (Guell et al. 2018), and cerebellar networks have been mapped even at the individual level (Marek et al. 2018). These diverse behavioral contributions and resting state networks make an understanding of the cerebellum in older adults important, particularly given that both behavioral domains are impacted in advanced age (e.g., Park et al. 2001; Seidler et al. 2010). Work over the past two decades has increasingly pointed to the cerebellum as a contributor to age-related performance differences when investigating volume (MacLullich et al. 2004; Bernard and Seidler 2013; Miller et al. 2013; Koppelmans et al. 2015, 2017), and we have known for quite some time that cerebellar volume is smaller in older adults and declines over time (Raz et al. 1998, 2005, 2010, 2013; Han et al. 2020). Furthermore, there are age differences in lobular cerebellar connectivity as well (Bernard et al. 2013), and subcortically, differing patterns of connectivity between the cerebellum and basal ganglia (Hausman et al. 2020).

Within the cerebellum are a series of nuclei, which are the primary output nodes of the structure. The largest of these nuclei, the dentate, provides the primary output to the cerebral cortex. Work in non-human primates has demonstrated that there are two dissociable circuits connecting the dentate with the cerebral cortex, via the thalamus. The more dorsal aspects of the dentate are associated with motor cortical regions, while the ventral aspect of the dentate is connected with the lateral prefrontal cortex (Dum and Strick 2003a). In the human brain, we used resting state functional connectivity (fcMRI) to investigate these dissociable circuits (Bernard et al. 2014). We used cerebellar-specific normalization methods (Diedrichsen 2006; Diedrichsen et al. 2009) in conjunction with single-voxel seeds, and we demonstrated that the dorsal and ventral dentate networks can in fact be dissociated in the human brain using fcMRI (Bernard et al. 2014). This dissociation has been further confirmed using high-resolution diffusion tensor imaging in the human brain (Steele et al. 2017).

Though our understanding of cerebellar connectivity, particularly that of the cerebellar dentate nucleus has greatly expanded in recent years, it remains unknown whether networks of the dentate nucleus are different across the lifespan. That is, there may be age relationships wherein connectivity is lower as individuals get older. However, it is also possible that in advanced age, these networks include different cortical regions as well. While prior work has demonstrated that connectivity of the cerebellar lobules is lower in OA (Bernard et al. 2013; Hausman et al. 2020), to date, the dentate nucleus has not been investigated in a sample encompassing the complete adult lifespan. While cerebellar lobular approaches are informative and reflect purported differences in processing and communication with the cerebral cortex (Bernard et al. 2013), given that the dentate nucleus is a critical aspect of this circuit as the primary output region to the cerebral cortex, further investigation is warranted. As such, we used a large data set including individuals across adulthood from ages 18 to 88 available from the Cambridge Center for Ageing Neuroscience (CamCAN) repository (Shafto et al. 2014; Taylor et al. 2017). We tested the hypothesis that dentate connectivity is impacted with advanced age and predicted that connectivity of both the dorsal and ventral dentate would be lower with increased age. In parallel, we were also interested in whether our initial findings using cerebellar specific methods would replicate in a large sample such as this, using more general analysis approaches. That is, can we still effectively dissociate dorsal and ventral dentate networks in the human brain with standard imaging analysis pipelines? By this, we refer to pipelines implemented in commonly available imaging software, such as the CONN toolbox (Whitfield-Gabrieli and Nieto Castañón 2012), without separate normalization of the cerebellum and the use of associated cerebellar toolboxes (SUIT; Diedrichsen et al. 2009). We recognize, however, that at this point there is no single common imaging pipeline used across the field, or one specific gold standard.

In advanced age females are more likely than males to develop Alzheimer’s disease (Carter et al. 2012; Mazure and Swendsen 2016), they are at a greater risk of falls (Stevens and Sogolow 2005; Hartholt et al. 2011), and suffer from frailty to a higher degree than their male counterparts (Puts et al. 2005). As such, an understanding of the brain in males and females is particularly important for future work seeking to understand and mitigate these sex differences in late life health outcomes.

When looking at associations between age and cerebellar metrics, to this point, investigations of sex differences are relatively limited, despite evidence to suggest that there may be sex differences in cerebellar structure (Raz et al. 1998; Bernard et al. 2015a; Han et al. 2020). In our own work investigating lobular cerebellar volume across adolescence through middle age we found that associations between regional volume and age differed in males and females. In the posterior cerebellum, females were best fit using a quadratic function such that volumes were largest in those in their late 20s and 30s, and there was a sharp decrease in volume in early middle age (Bernard et al., 2015). This is in contrast to males that showed linear relationships with age. More generally, Steele and Chakravarty (Steele and Chakravarty 2018) demonstrated sex differences in adults in lobular volume. Critically however, though females live longer than males, they experience poorer later life outcomes, as noted above. While falls are no doubt compounded by peripheral changes in musculature and the incidence of osteoporosis, health outcomes in older females underscore the importance of detailed investigation into factors that may contribute to these trajectories. Given that the cerebellum has been implicated in a variety of motor and cognitive domains (e.g., Stoodley et al. 2012; King et al. 2019), and more recently has been implicated in Alzheimer’s disease (Tabatabaei-Jafari et al. 2017; Jacobs et al. 2018; Toniolo et al. 2018; Olivito et al. 2020), understanding sex differences in dentate connectivity with age will provide important new insights into sex differences in cerebellar connectivity in adulthood. Though literature in this area is relatively limited, we hypothesized that females would show greater effects of age such that there are more extensive negative correlations between age and dentate connectivity in both the dorsal and ventral dentate, relative to males.

## Methods

### Data

Data used in the preparation of this work were obtained from the CamCAN repository (available at http://www.mrc-cbu.cam.ac.uk/datasets/camcan/)(Shafto et al. 2014; Taylor et al. 2017). Participants included in this investigation had both resting state and structural brain imaging data available. After excluding individuals with missing resting state data, the final sample included 590 people (297 females) between the ages of 18 and 88 (mean age 54.54 ± 18.65 years; females: 18-88, mean 53.82 ± 18.94 years; males: 18-87, mean 55.31 ± 18.38 years). Data were collected using a 3T Siemens TimTrio. Complete data collection details can be found online (see link above), and are outlined by Shafto and colleagues (2014) and Taylor and colleagues (2017). The former also includes data on participant sampling approaches. For our analyses here, we used the T1 MPRAGE along with the resting state EPI scans. The resting state scan was approximately 8:30 minutes long and was completed in one session with a TR of 1.97 seconds. The voxel size of the acquired resting state data was 3×3×4.4mm. Raw data were acquired and used here.

### Processing and Analysis

Methods for data processing used here parallel those from our recent work (Bernard et al. 2017; Hausman et al. 2020) and have been reported here for the sake of completeness and transparency. All data were processed and analyzed using the CONN toolbox version 19b (Whitfield-Gabrieli and Nieto Castañón 2012) implemented in conjunction with Matlab 2019b. All analyses were conducted with the advanced computing resources provided by Texas A&M High Performance Research Computing. We followed the standard preprocessing pipeline in CONN, including functional realignment and unwarping, functional centering of the image to (0, 0, 0) coordinates, slice-timing correction, structural centering to (0, 0, 0) coordinates, structural segmentation and normalization to MNI space, functional normalization to MNI space, and spatial smoothing with a smoothing kernel of 5mm FWHM. This paralleled the approach taken in recent investigations conducted by our group (Bernard et al. 2017; Hausman et al. 2020). Because of the potential confounding effects of motion and signal outliers (Power et al. 2012; Van Dijk et al. 2012) these procedures also included processing using the Artifact Rejection Toolbox (ART). This was set using the 95^th^ percentile settings and allowed for the quantification of participant motion in the scanner and the identification of outliers based on the mean signal. These effects, as well as white matter and cerebral-spinal fluid signals, were included as confounds and regressed out during denoising, prior to first level analysis. Data were denoised using a band-pass filter of .008 – .09 Hz. With these settings, the global-signal z-value threshold was set at 3, while the subject-motion threshold was set at .5 mm. 6-axis motion information and frame-wise outliers were included as covariates in our subsequent first-level analyses. Notably however, these frame-wise outliers are not removed, but the signal is de-spiked to bring it closer to the global mean during the denoising process. The 6 motion covariates were reduced to one variable by averaging the absolute value of each axes’ average and the frame-wise time-series was averaged across each participant, resulting in one value for each measure for each participant for group comparison.

Seeds used for analysis were placed in both the dorsal and ventral dentate using regions defined in our earlier work investigating dissociable human dentate circuits (Bernard et al. 2014). The dorsal dentate seed was centered on (12, −57, −30) while the ventral seed was centered on (17, −65, −35) (Figure 1). Seeds were placed in the right dentate, as this corresponds to the dominant motor hemisphere for right-handed individuals, such as those included here. We were particularly interested in the dominant motor hemisphere for consistency with past work. Further, given the analyses of sex in addition to the associations with age, we were cognizant of limiting the number of comparisons and as such, limited our seeds to one hemisphere. Spherical seeds corresponding in size to a single voxel (3mm diameter) were created and used in all analyses. While recent work has suggested there may be a third sub-region in the dentate nucleus (Guell et al. 2020), we focused here on only two sub-regions. Primarily, this is due to the known anatomical underpinnings of these particular circuits. While Guell and colleagues used a parcellation approach with resting state data (2020), to this point, there are no known white matter circuits subserving a third region. While future work may resolve additional white matter tracts in the non-human and human primate brains, our aim here was to stick with functional networks known to map on to underlying structural circuitry. In addition, given the voxel size of the images available through the CamCAN repository, our goal was to eliminate overlap between the seed territories, and as such we limited ourselves to these two regions.

**Figure 1.**
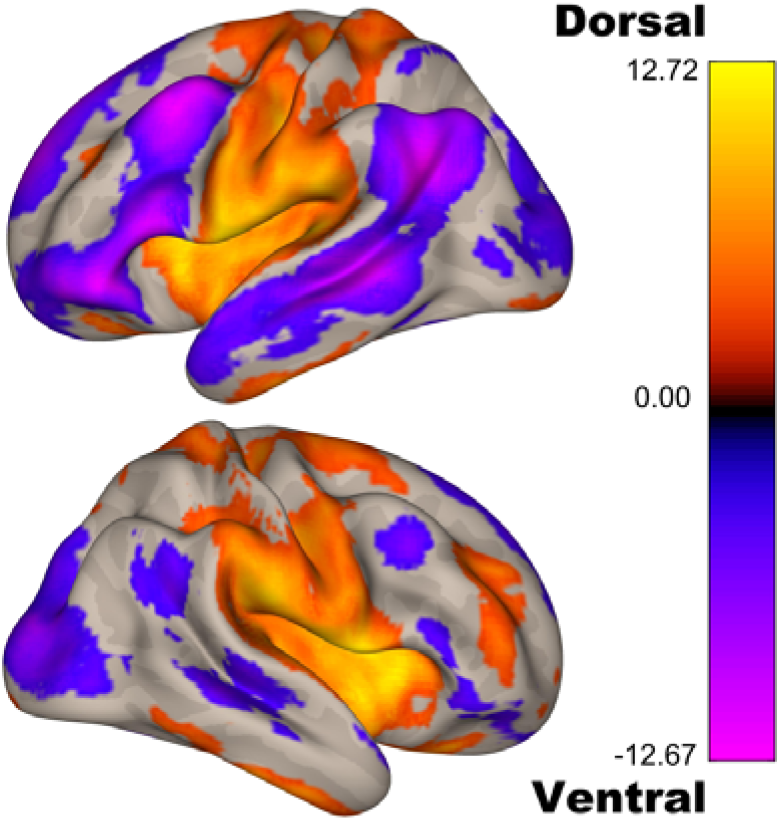
Dorsal and ventral dentate seeds locations. These seeds parallel those used by Bernard and colleagues (2014) and were chosen to be non-overlapping as pictured here.

After pre-processing we completed first-level and group-level analysis, also using the CONN toolbox. At the whole group level, we first replicated our prior analyses investigating dorsal and ventral dentate connectivity using standard imaging analysis approaches. First, we delineated the connectivity patterns for each seed independently and conducted group-level analyses to produce connectivity maps for the dorsal and ventral dentate. To further understand the dissociability of these networks, we ran a statistical contrast between the two seeds (dorsal>ventral; ventral>dorsal), and we computed a semi-partial correlation such that when looking at the dorsal seed we controlled for signal from the ventral and vice versa. Next, we conducted a regression with participant age to determine the relationship between age and dentate network connectivity. Our primary analyses were a standard bivariate correlation, though we also included exploratory quadratic analyses. We conducted these analyses separately for the dorsal and ventral seed. Finally, we investigated whether and how these patterns may differ in males and females across adulthood. We conducted correlation analyses between age and dentate seed connectivity in males and females separately (both linear and quadratic). Further, we directly compared dentate connectivity patterns between the two sexes and investigated interactions in the relationships with age and sex. In all cases, analyses were evaluated using a voxel threshold of p<.001, followed by a cluster threshold that was FDR corrected at p<.05.

## Results

As noted above, all data was processed with head motion and image outliers in mind, so that these could be used in the subsequent connectivity analyses. Data about motion, including mean motion (mm) as well as maximum motion (mm), are both included in Supplementary Table 1, where in participants have been grouped based on age, in decades. Perhaps not surprisingly, given that existing work has demonstrated increases in motion with increasing age (e.g., Saccà et al. 2021), we too saw significant positive correlations with age for both mean motion (r_(590)_=.469, p<.01) and maximum motion (r_(590)_=.245, p<.001). Furthermore, when comparing age groups (by decade) there is also a significant main effect of age for both mean (F_(6,583)_=28.40, p<.001) and maximum (F_(6,583)_=8.32, p<.001) motion. With that said, as outlined above, we included motion variables in all of our analyses, and outliers were de-spiked. As such, despite these associations with age, given the motion covariates that were included in the resting state analyses, we believe that these results provide important insights into dentate connectivity in the aging brain.

### Dorsal and Ventral Networks

First, across the whole sample we investigated the dorsal and ventral dentate networks. While we have previously shown a dissociation between the dorsal and ventral networks in a small sample using cerebellar-specific methods (Bernard et al. 2014), our findings here suggest that these networks can be dissociated across large representative adult samples using more traditional brain imaging analysis approaches (that is, analyses designed for the whole-brain, using readily available imaging toolboxes). First, the dorsal dentate seed shows robust connectivity with primary and premotor areas bilaterally (Figure 2, Supplementary Table 2). Notably however in the right hemisphere, there were also areas of correlation with the lateral prefrontal cortex. Second, the ventral dentate showed robust connectivity with the prefrontal cortex, parietal, and temporal cortices (Figure 2; Supplementary Table 2). These findings are consistent with our prior work (Bernard et al. 2014), and other replications (Steele et al. 2017; Guell et al. 2020). Statistical comparisons between the two seeds also support the two distinct dentate networks (Supplementary Table 2).

**Figure 2.**
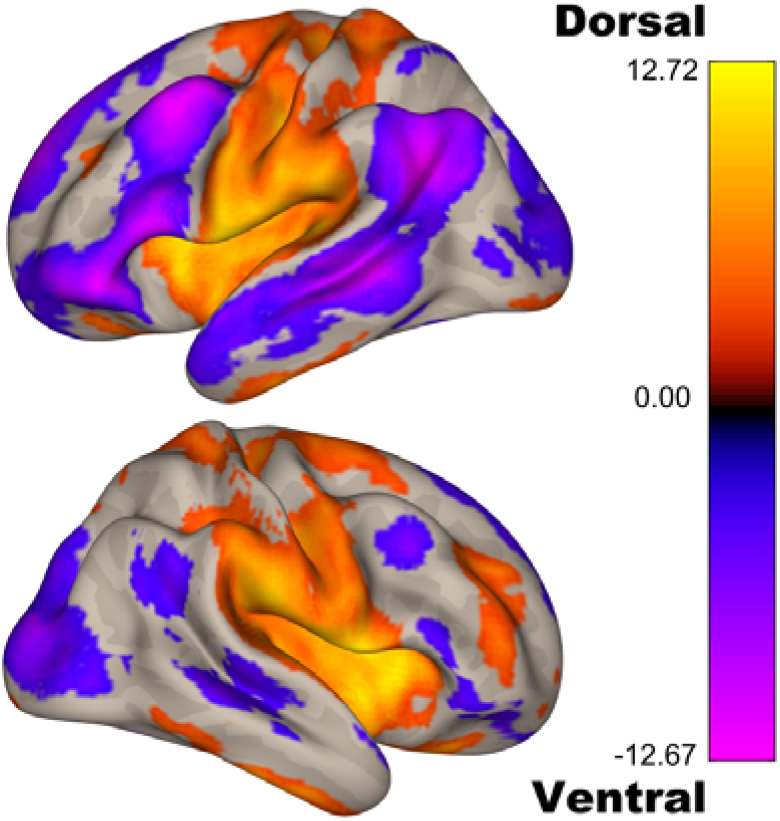
Functional connectivity patterns for the dorsal (yellow/orange) and ventral (blue/purple/pink) dentate seeds. The patterns of connectivity for the two seeds parallel the dissociation seen in non-human primates and in human work, even when using traditional whole-brain processing and analysis approaches.

To further investigate the dissociability of these two networks, we also analyzed the data using a semi-partial correlation approach, which has previously been implemented in other brain regions with seeds that are close in space (Adams et al. 2019). The results of this analysis are presented in Supplementary Figure 1 and Supplementary Table 3. In brief, we see a similar dissociation between the two seeds when controlling for the signal in the other seed, providing more support for the distinct patterns of connectivity of the dorsal and ventral dentate in the human brain.

### Associations With Age Across Adulthood

After replicating the dorsal and ventral dentate connectivity dissociation in this large adult sample, we investigated the age-associations for each seed. When investigating the dorsal dentate, we found that connectivity does in fact get lower with increasing age in pre-motor, motor, and somatosensory regions of the network (Figure 3, Table 1). Investigations of the ventral dentate nucleus seed primarily revealed negative correlations with age. With increasing age, connectivity is lower in the parietal, and anterior temporal lobe regions, as well as the frontal pole and putamen. Somewhat surprisingly however, there was a positive correlation with the anterior cingulate such that as age increased, so did the correlation with the signal in the ventral dentate nucleus. Notably, we did not use a limiting mask with the originally defined networks so as to fully explore these relationships with age, and as such some of the areas are outside the initially defined networks.

**Table 1.**
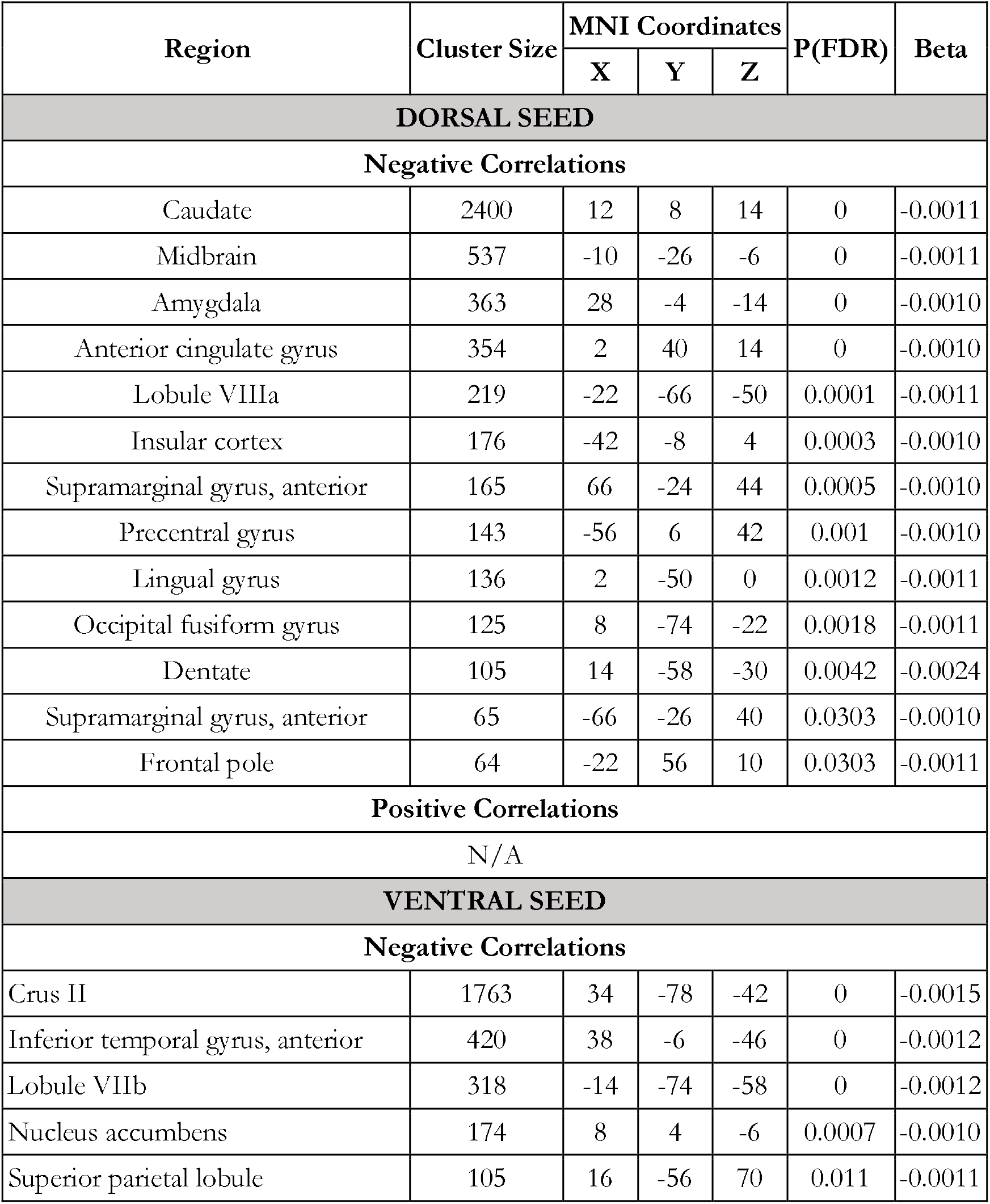
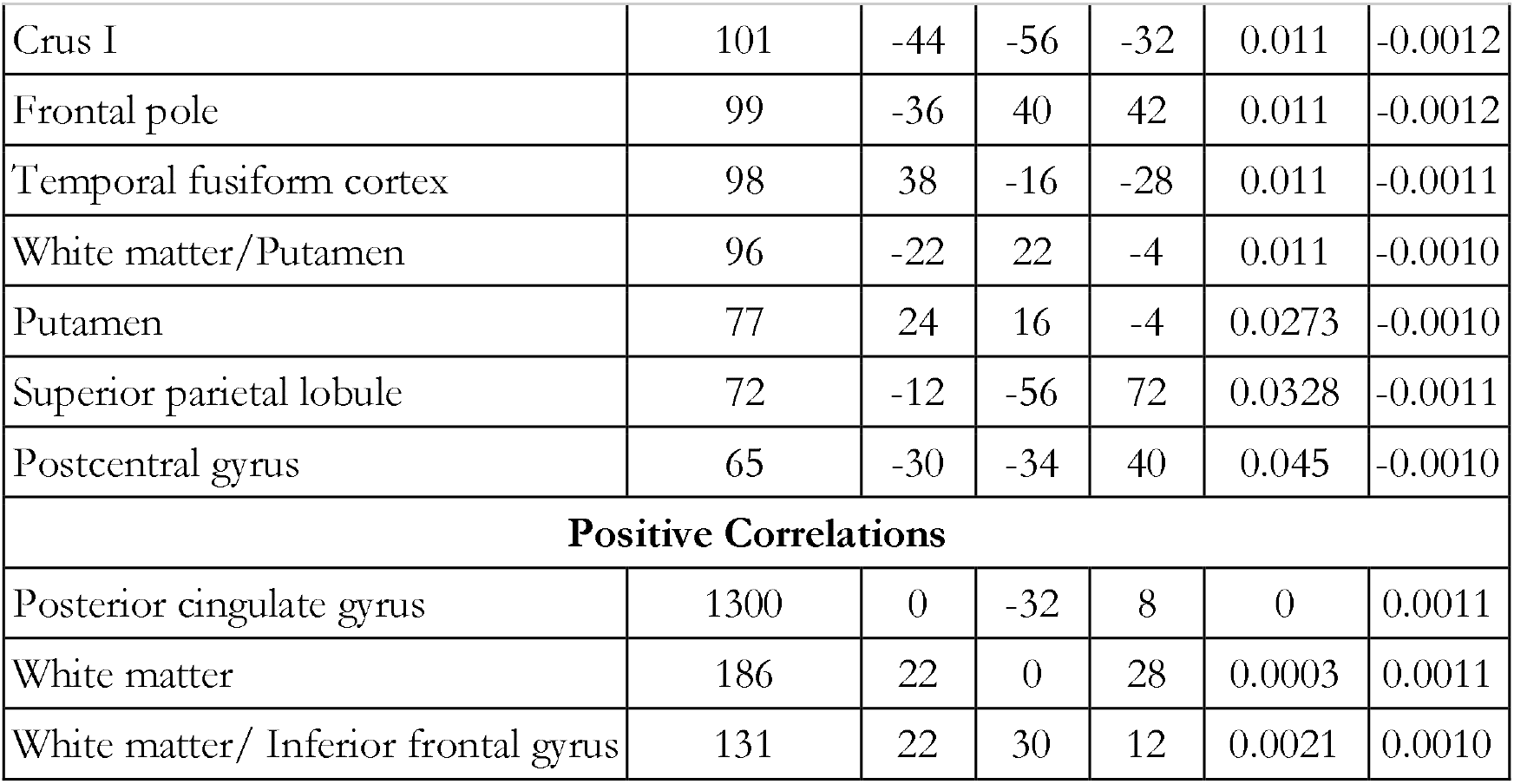
Dentate Connectivity Across the Adult Lifespan. Positive and negative correlations with age for both the dorsal and ventral dentate nucleus seeds. Anatomical locations were determined using the Harvard-Oxford max probability atlas while subcortical regions were localized with the JHU atlas in MRICron. Cerebellar locations were determined using the SUIT atlas. Beta values are unstandardized.

**Figure 3.**
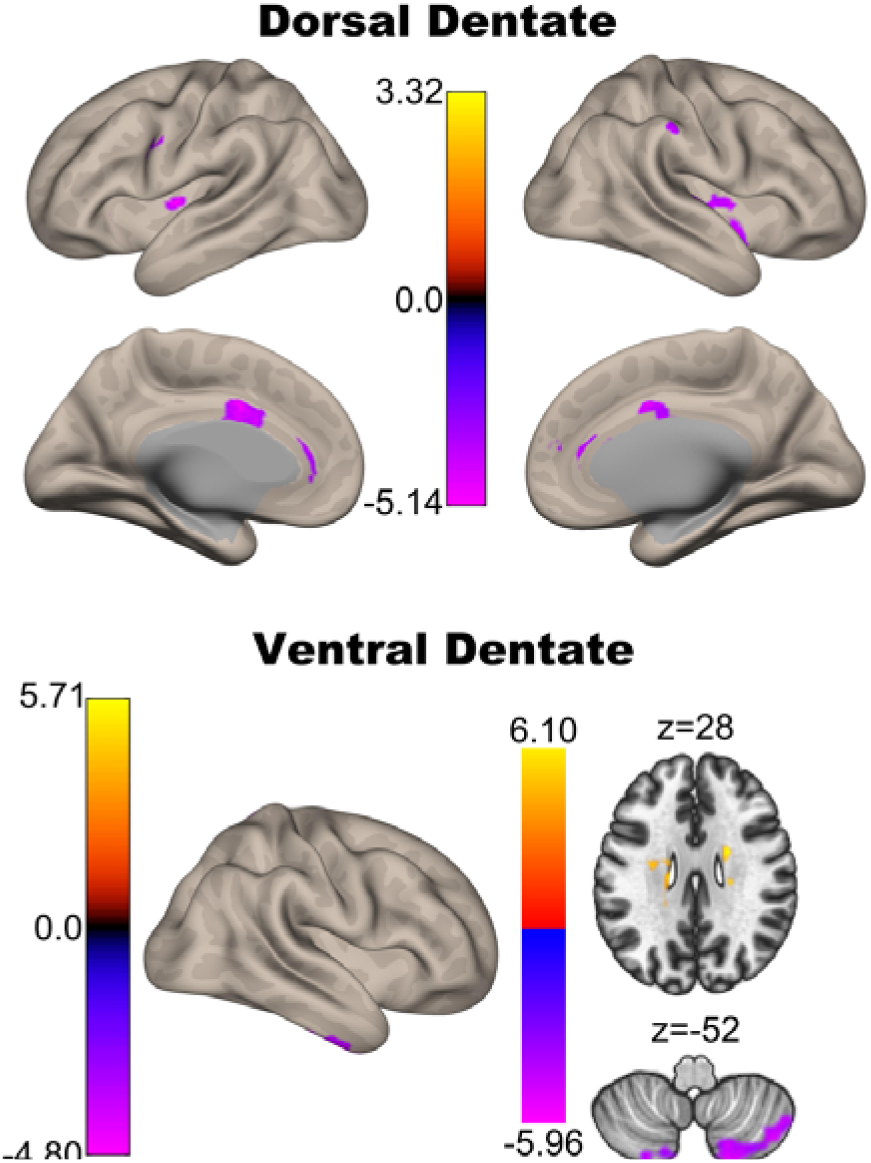
Associations between connectivity and age for the dorsal (top) and ventral (bottom) dentate seeds. While there are some positive associations reported (see Table 1), the views here primarily demonstrate negative associations between connectivity strength and age (purple), though some small regions of positive correlation (yellow/orange) with the ventral dentate seen are visible in the temporal lobe in the axial slice.

As work investigating lobular volume with age has demonstrated some quadratic relationships particularly when investigating females and males separately (Bernard et al. 2015b; Han et al. 2020), we also conducted exploratory quadratic analyses. When looking at the entire sample, quadratic relationships were only revealed for the dorsal dentate seed (Supplementary Table 4). There were positive quadratic relationships (an “inverted-u” pattern) that were constrained to the cerebellum in Lobule V and Vermis VI, both areas that are associated with motor performance and networks (Dum and Strick 2003a; Stoodley et al. 2012; King, Hernandez-castillo, Sereno, et al. 2019). Additionally, there were negative quadratic relationships (“u-shaped” pattern) with the paracingulate, precuneus, and frontal orbital cortex. There were no significant relationships when investigating the ventral seed.

### Sex Differences

Finally, we completed analyses of sex differences in cerebellar dentate connectivity. A growing literature indicates sexual dimorphism of the cerebellum that is present in older adulthood (e.g., Raz et al. 1998; Bernard et al. 2015; Han et al. 2020). While the literature to date has primarily focused on cerebellar structure and volume, here we investigated sex differences in dentate connectivity when collapsing across the entire sample as well as with respect to age in both males and females. When collapsing across the sample to look at sex differences in network connectivity, several differences were observed. Males showed greater connectivity relative to females between the dorsal dentate and Lobule VI, superior frontal gyrus, temporal pole, and occipital pole. For ventral dentate males showed greater connectivity than females with the temporal occipital fusiform cortex, and the parahippocampal gyrus. Females showed greater connectivity relative to males between the dorsal dentate and temporal lobe regions as well as Lobule VIIIa (Table 2). There were no areas where ventral dentate connectivity was greater for females relative to males. Thus, in this sample, collapsing across the adult lifespan, there are some sex differences in connectivity in the dorsal and ventral dentate networks..

**Table 2.**
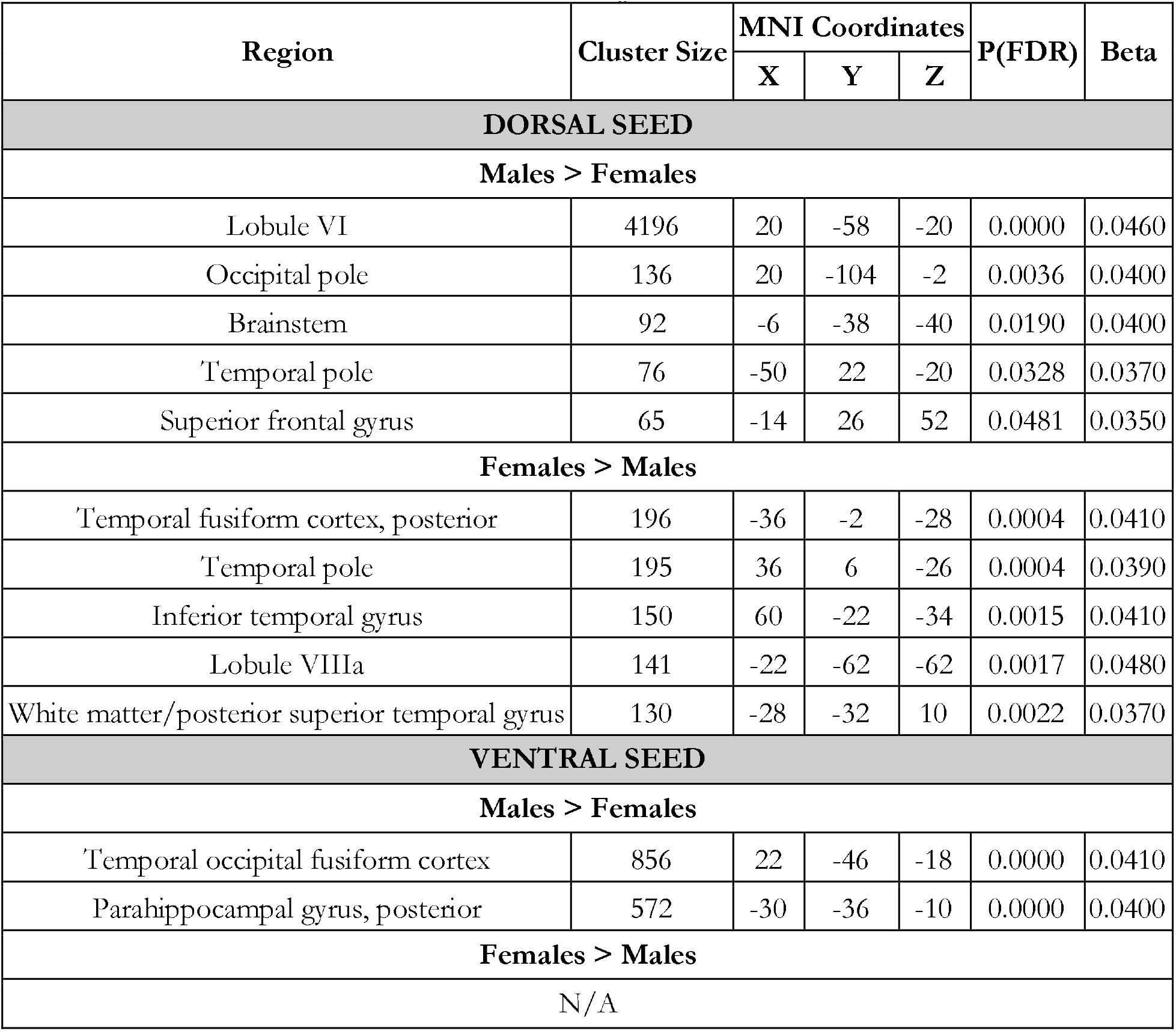
Sex Differences in Dentate Connectivity. Anatomical locations were determined using the Harvard-Oxford max probability atlas and subcortical areas were localized with the JHU atlas in MRICron. Cerebellar locations were determined using the SUIT atlas. Betas are unstandardized.

There are also notable connectivity associations with age in both males and females. With increasing age in females, there is significantly lower connectivity between the dorsal dentate and premotor cortical regions (superior frontal gyrus), frontal pole, brain stem and midbrain, and the anterior cingulate cortex. In the ventral dentate, connectivity is lower in older females again in the anterior cingulate, the superior parietal lobule, and the supramarginal gyrus. For both seeds there were some positive, potentially spurious, correlations that are primarily localized to white matter. Males, however, demonstrated distinct correlations with different regions, compared to females. Older males show lower connectivity between the dorsal dentate and the supramarginal gyrus, caudate and thalamus, while with the ventral dentate there are negative correlations with age and connectivity in the temporal lobe (inferior and middle temporal gyrus, temporal fusiform), and lobule VIIb. The only positive correlation in males was with the ventral dentate and posterior cingulate cortex. Detailed findings are presented in Tables 3 and 4 and are visualized in Figure 4. For both males and females, we also investigated quadratic relationships. In general, there were minimal quadratic associations, and these were more prominent in males, though localized to regions within the cerebellum (Supplementary Tables 5 and 6).

**Table 3.** Dorsal and Ventral Dentate Connectivity Associations with Age in Females. Positive and negative correlations with age for both the dorsal and ventral dentate nucleus seeds. Anatomical locations were determined using the Harvard-Oxford max probability atlas and subcortical regions were localized using the JHU atlas in MRICron. Cerebellar locations were determined using the SUIT atlas. Beta values are unstandardized.

| Region | Cluster Size | MNI Coordinates |  |  | P(FDR) | Beta |
| --- | --- | --- | --- | --- | --- | --- |
|  |  | X | Y | Z |  |  |
| DORSAL SEED |  |  |  |  |  |  |
| Negative Correlations |  |  |  |  |  |  |
| Brainstem | 811 | 12 | -16 | -14 | 0 | -0.0014 |
| Anterior cingulate gyrus | 391 | 8 | 34 | 8 | 0 | -0.0014 |
| Dentate | 136 | 14 | -56 | -32 | 0.0022 | -0.0032 |
| Frontal orbital cortex | 121 | -10 | 16 | 26 | 0.0034 | -0.0014 |
| Frontal pole | 105 | 22 | 38 | 42 | 0.0061 | -0.0014 |
| White matter/globus pallidus | 93 | -10 | 4 | 2 | 0.0095 | -0.0014 |
| Dentate | 89 | -14 | -56 | -36 | 0.0101 | -0.0018 |
| Superior frontal gyrus | 60 | -22 | 10 | 56 | 0.0473 | -0.0013 |
| Positive Correlations |  |  |  |  |  |  |
| Inferior temporal gyrus | 144 | -40 | -46 | -4 | 0.002 | 0.0014 |
| White matter/caudate | 97 | 20 | 4 | 26 | 0.0064 | 0.0014 |
| White matter | 96 | -24 | -12 | 30 | 0.0064 | 0.0013 |
| White matter/caudate | 93 | 18 | 28 | 2 | 0.0064 | 0.0015 |
| Middle temporal gyrus | 62 | 40 | -48 | -2 | 0.0297 | 0.0014 |
| White matter | 54 | -30 | -24 | 30 | 0.0409 | 0.0014 |
| VENTRAL SEED |  |  |  |  |  |  |
| Negative Correlations |  |  |  |  |  |  |
| Superior parietal lobule | 124 | 14 | -54 | 72 | 0.0119 | -0.0015 |
| Dentate | 100 | 16 | -66 | -36 | 0.0198 | -0.0034 |
| Stria terminalis | 91 | -26 | -30 | 4 | 0.0212 | -0.0014 |
| Anterior cingulate gyrus | 75 | -2 | 4 | 42 | 0.0384 | -0.0015 |
| Supramarginal Gyrus, anterior | 69 | 68 | -26 | 36 | 0.0435 | -0.0013 |
| Positive Correlations |  |  |  |  |  |  |
| Precuneus | 300 | 20 | -44 | 20 | 0 | 0.0015 |
| White matter | 106 | -18 | -16 | 30 | 0.0063 | 0.0014 |
| White matter | 66 | 22 | 0 | 28 | 0.0377 | 0.0014 |

**Table 4.** Dorsal and Ventral Dentate Connectivity Associations with Age in males. Positive and negative correlations with age for both the dorsal and ventral dentate nucleus seeds. Anatomical locations were determined using the Harvard-Oxford max probability atlas and subcortical regions were localized using the JHU atlas in MRICron. Cerebellar locations were determined using the SUIT atlas. Beta values are unstandardized.

| Region | Cluster Size | MNI Coordinates |  |  | P(FDR) | Beta |
| --- | --- | --- | --- | --- | --- | --- |
|  |  | X | Y | Z |  |  |
| DORSAL SEED |  |  |  |  |  |  |
| Negative Correlations |  |  |  |  |  |  |
| Caudate | 397 | -10 | 4 | 12 | 0 | -0.0014 |
| Thalamus | 209 | 20 | -20 | 18 | 0.0002 | -0.0015 |
| Supramarginal gyrus, anterior | 128 | 64 | -26 | 44 | 0.0039 | -0.0014 |
| Thalamus | 108 | -18 | -22 | 18 | 0.0078 | -0.0015 |
| Positive Correlations |  |  |  |  |  |  |
| N/A |  |  |  |  |  |  |
| VENTRAL SEED |  |  |  |  |  |  |
| Negative Correlations |  |  |  |  |  |  |
| Lobule VIIb | 1575 | 36 | -68 | -54 | 0 | -0.0019 |
| Inferior temporal gyrus, posterior | 325 | 44 | -12 | -42 | 0 | -0.0017 |
| Middle temporal gyrus, posterior | 231 | -46 | -24 | -10 | 0.0001 | -0.0016 |
| Temporal fusiform cortex, posterior | 110 | 38 | -16 | -30 | 0.0103 | -0.0016 |
| Positive Correlations |  |  |  |  |  |  |
| Posterior cingulate gyrus | 150 | -12 | -36 | 14 | 0.0035 | 0.0014 |

**Figure 4.**
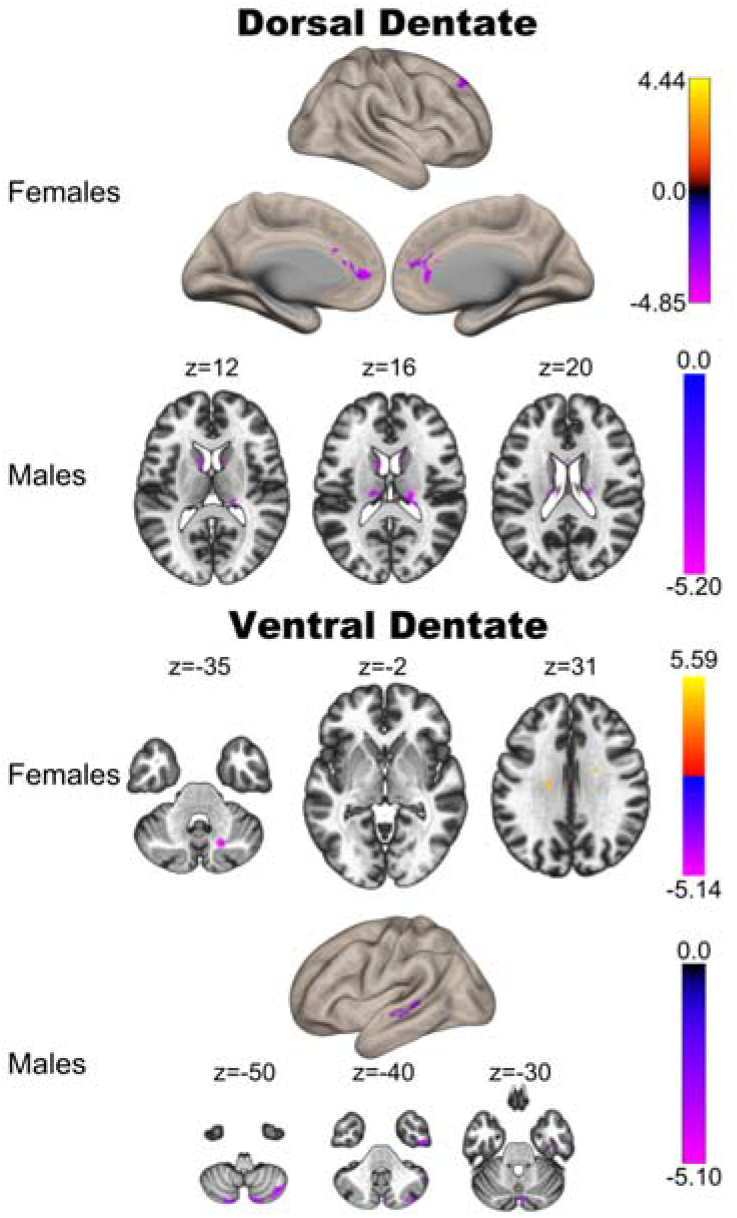
Correlations between the dorsal (top) and ventral (bottom) regions of the dentate nucleus with age in females (top) and males (bottom). Notably, the negative correlations are more extensive in females, particularly for the dorsal dentate seed.

Finally, we also investigated differences in the relationships between connectivity and age (Supplementary Table 7). Here, we found that in females, there was a stronger association between dorsal dentate connectivity with the parietal operculum and age, while in males this was the case for the angular gyrus. With respect to the ventral dentate, the age relationship with the posterior aspects of the superior temporal gyrus was larger in females. There were no regions where connectivity-ag relationships were stronger in males. Together, these results suggest that there are some sex differences in age relationships with respect to networks of the dorsal and ventral dentate nucleus.

## Discussion

Here, using a large sample of adults representing the adult lifespan, we investigated patterns of resting state connectivity in the dorsal and ventral dentate nucleus. First, we demonstrated that we can replicate prior results computed with cerebellar-specific analyses (that is, separate cerebellar normalization in a cerebellar atlas separated from the rest of the cerebral cortex) when using more standard imaging analysis protocols (e.g., the CONN toolbox, or other commonly used neuroimaging processing packages). We demonstrated a clear dissociation in the patterns of dorsal and ventral dentate connectivity, such that the dorsal dentate was robustly connected with primary and pre-motor cortical regions, while the ventral dentate was more strongly correlated with pre- frontal and association areas of the cerebral cortex. Furthermore, we demonstrated that across adulthood, connectivity between both seeds and their associated cortical networks was lower in advanced age. In older individuals, connectivity between dentate sub-regions is smaller. Finally, we demonstrated that in females and males there are distinct patterns of connectivity associations with age.

### Dentate Dissociation Replication

Work in non-human primates elegantly demonstrated that the dorsal and ventral dentate nucleus territories are part of parallel circuits with the motor and prefrontal regions of the cortex, respectively (Dum and Strick 2003b). Initial work seeking to map these same circuits functionally at rest in the human brain relied upon data processing approaches that were optimized for cerebellar imaging (Bernard et al. 2014). That is, the cerebellum was separated from the cortex, and normalized to a cerebellar-specific atlas, as opposed to using one processing stream with the whole brain, such as what is implemented in the CONN toolbox. While this work demonstrated that the human dentate nucleus does have at least two distinct territories with dissociable circuits in the human brain, whether or not these circuits can be dissociated using whole-brain methods was to this point, unknown. Here, we used traditional processing and whole-brain methods implemented in the CONN toolbox while investigating the dissociability of the dorsal and ventral dentate nucleus sub-regions. Consistent with our prior work, as well as subsequent work replicating this pattern of dissociation in the human brain (Steele et al. 2017; Guell et al. 2020), we further demonstrated that the dorsal dentate region is more robustly correlated with motor cortical regions, while the ventral dentate is more robustly correlated with prefrontal and association cortices. Together, this demonstrates that more general processing parameters may be sufficient for dissociating these networks, at least in larger samples, such as the one used here.

### Dentate Network Connectivity Across Adulthood

The primary goal of our work here was to determine whether networks of the cerebellar dentate nucleus follow patterns with age that parallel those of the cerebellar cortex. Across both seed regions, we saw robust negative correlations with age, consistent with our prior work demonstrating lower cerebellar connectivity in advanced age (Bernard et al. 2013; Hausman et al. 2020). Notably, these negative correlations with age correspond to the key areas of the dorsal and ventral dentate networks themselves. That is, in the dorsal dentate, in older individuals, connectivity is lower in primary motor and somatosensory regions, while in the ventral dentate connectivity is lower in frontal, parietal, and temporal lobe regions. More broadly, this is consistent with widespread age differences in resting state network dynamics within networks as individuals age, seen across cortical networks (Andrews-Hanna et al. 2007; Tomasi and Volkow 2012; Ferreira and Busatto 2013; Ferreira et al. 2016). This is also consistent with work showing lower connectivity of the cerebellar lobules in OA (Bernard et al. 2013). Notably however, for both seeds there were associations with age outside the dorsal and ventral dentate seeds themselves, suggesting that there may also be differences in between-network interactions in OA. Further, longitudinal evidence suggests that there are in fact changes over time in cortical resting state connectivity in OA (Oschmann and Gawryluk 2020). However, it is also critical to note the multi-directionality in these results. While there are negative associations between connectivity and age in the core areas associated with the dorsal and ventral dentate networks, the ventral seed also showed positive correlations with brain regions outside the core networks themselves. Though somewhat surprising, these connectivity increases may be reflective of an attempt at compensation or use of neural scaffolding to maintain function (Park and Reuter-Lorenz 2009; Reuter-Lorenz and Park 2014; Cabeza et al. 2018). Alternatively, this may be indicative of a decreased efficiency in brain network organization. However, without behavioral data and associations we can only speculate.

Critically, these results provide further insights as to the extent to which cerebellar networks are impacted in older adulthood. Perhaps not surprisingly, the cerebellar dentate nucleus is also negatively impacted in advanced age, as quantified by resting state connectivity. There are two key considerations for these findings. First, there is an open question as to the mechanism driving these negative relationships with age. That is, why do OA show lower cerebellar connectivity, relative to young? With respect to the cerebellum specifically, we and others have previously demonstrated age differences in lobular volume (Bernard and Seidler 2013; Koppelmans et al. 2015, 2017; Han et al. 2020), and more recent investigations have confirmed these differences and also noted volumetric declines over time (Han et al. 2020). We have speculated that differences in volume as well as connectivity may negatively impact cerebellar processing in advanced age (Bernard and Seidler 2014), which in turn may contribute to the performance differences seen across the motor and cognitive domains in OA. Here however, we have extended findings to indicate that dentate connectivity is also negatively impacted in aging. With that in mind, given that these nuclei serve as the output region of the cerebellum and receive input from the cerebellar cortex, the age differences in dentate connectivity reported here may reflect differences in lobular processing. Investigating the relative associations between lobular and dentate connectivity in the context of behavior and functional outcomes in the future may begin to tease apart the respective contributions of age differences in lobular and dentate connectivity.

However, changes in neurotransmitter systems may also contribute to the age differences in dentate connectivity. Previously, we have demonstrated marked differences in connectivity between the cerebellum and basal ganglia in OA relative to YA (Bernard et al. 2013; Hausman et al. 2020) using lobular cerebellar seeds. We speculated that age differences in dopamine may be contributing, at least in part, to the differences in connectivity in advanced age. As noted, work by Kelly and colleagues (Kelly et al. 2009) demonstrated that the administration of l-dopa increased connectivity between the striatum and cerebellum in healthy YA. Here, though we are looking at dentate connectivity, we suggest that normative age-differences in dopamine (McGeer et al. 1977; Fearnley and Lees 1991) may be contributing in part to the differences seen here. In parallel, other neurotransmitter differences in advanced age may also be impacting cerebellar connectivity patterns. Serotonin, GABA, and acetylcholine all act in the cerebellum (Oostland and van Hooft 2016) and are impacted in advanced age. Normative age-differences in neurotransmitter action may in turn impact cerebellar processing (as well as cerebral cortical processing) and associated connectivity patterns at rest. Targeting these neurotransmitters, in conjunction with investigations of connectivity in the future stands to provide new insights into these age differences.

The second area of consideration for these findings is with respect to their implications for our understanding of age-related disease, particularly mild cognitive impairment and dementia. Though historically the cerebellum has not been a major target of investigation in dementia, and more specifically in Alzheimer’s Disease (AD), in recent years, this has changed (for a review see Jacobs et al. 2018). Notably, converging evidence indicates that the cerebellum is impacted in AD. Cerebellar decline has been linked to functional decline in advanced AD (Tabatabaei-Jafari et al. 2017), and there is additional evidence to indicate that there are relationships between the cerebellar volume and cognition in mild cognitive impairment (Lin et al. 2020). In a recent meta-analysis Gellersen and colleagues (Gellersen et al. 2020) demonstrated that in the cerebellum, there are some overlapping areas of cerebellar structural loss when comparing AD to normative healthy aging. However, differences were also observed between the two groups wherein atrophy in AD was primarily lateralized to the right hemisphere (Gellersen et al. 2020). Broadly, this work suggests an increasing need for further investigation of the cerebellum in AD, as it may not be entirely spared structurally. With respect to functional networks, in patients with AD, cerebellar connectivity with the cortex is decreased (Zheng et al. 2017), and patterns of atrophy in the cerebellum are in areas associated with the default mode and salience networks, perhaps impacting general network coherence (Guo et al. 2016). Further, the dentate itself has been investigated in AD. Interestingly, Olivito and colleagues (2020) found *increased* connectivity between the dentate and regions of the medial temporal lobe in patients with AD relative to controls. However, they did not investigate the distinct dentate sub-regions (Olivito et al. 2020), which may have impacted their findings. This is somewhat surprising, given work on the cerebellar cortex (Zheng et al. 2017). We suggest that the work presented here represents an important point of comparison for that in clinical populations, particularly dementia, as we have characterized patterns of dentate connectivity across the adult lifespan with both the dorsal and ventral regions dissociated. Additional follow-up work comparing AD and mild cognitive impairment samples to this work in healthy adults will provide further insights into diverging trajectories in healthy aging and disease.

### Sex Differences in Dentate Connectivity

In addition to quantifying the associations between dorsal and ventral dentate nucleus connectivity and age across adulthood, we also investigated sex differences in dentate connectivity. Our results support sex differences in dentate connectivity, and notably, our results suggest that associations between dentate connectivity and age differ in females and males to some degree. This suggests potential sex differences in the process of aging with respect to cerebellar functional networks. Sex differences in cerebellar structure have been reported (Oguro et al. 1998; Raz et al. 2001; Tiemeier et al. 2010; Bernard et al. 2015a; Steele and Chakravarty 2017), though the results have been somewhat mixed to this point, particularly when looking at broad anatomical swaths of the cerebellum, such as an entire hemisphere, or the cerebellar vermis in its entirety. These different patterns however, are consistent with our own work showing sex differences in age-volume relationships when investigating regional cerebellar volume across adolescence through late middle age (Bernard et al. 2015a). While these differences were in volume of the cerebellar lobules, it suggests that cerebellar relationships with age may differ between the sexes as we see here.

With respect to fcMRI, sex differences in connectivity patterns have been previously reported (e.g., Biswal et al. 2010; Tomasi and Volkow 2012; Alaerts et al. 2016; Engman et al. 2016; Weis et al. 2019), though the direction of these differences is somewhat variable across studies and samples. In addition, in studies of aging sex differences have also been reported. Tomasi and Volkow (2012) found that in females default mode network connectivity was higher than males, but it was weaker in somatosensory networks. Scheinost and colleagues (Scheinost et al. 2015) looked across multiple cortical and subcortical networks and also found age by sex interactions, though the patterns of the interaction vary across networks. While there is converging evidence to indicate that there are sex differences in functional networks, particularly in the cortex, work on cerebellar networks is relatively limited to this point, and has been focused on clinical groups that show sex differences in prevalence, such as Autism spectrum disorder (e.g., Smith et al. 2019). Thus, our findings here are novel in that they provide insights into sex differences in cerebellar dentate connectivity when looking at the whole sample, and with respect to sex differences in the relationships between connectivity and age.

### Limitations

This investigation has provided important new insights into cerebellar dentate connectivity across the lifespan. However, this work is not without limitations. First, as we relied up on a publicly available dataset, we were unable to investigate targeted behaviors of interest, and the data collection parameters were outside of our control. Regarding the former, without targeted behavioral assessments, we are unable to make any inferences as to the functional implications of these age-associations with connectivity. Further this makes it especially challenging to understand the increases in connectivity with age. We have suggested two possibilities; that this may be a compensatory increase or indicative of decreased network efficiency. Behavioral associations would provide some insight as to the role of these increases in connectivity with age. The inclusion of behavioral metrics in conjunction with resting state data will be critical for future work looking to better understand the impact of these age differences in the brain with respect to behavior. Relatedly, the imaging data was collected using a set of parameters optimized for whole-brain imaging, with a voxel size slightly larger than what we would have aimed to use in an investigation targeting the dentate nucleus. Though prior work looking at lobular cerebellar connectivity in advanced age has suggested that volume is not associated with lower connectivity in older adults (Bernard et al. 2013), we were unable to investigate dentate volume here, given the small size and sub-region approach, and the imaging parameters. Thus, while evidence from prior work suggests this is likely not a key factor in the associations with age reported here, we cannot rule it out. With that said, the large sample represents an important advance relative to prior work, and critically, we replicated the dissociation in dorsal and ventral seed connectivity. This suggests that we were measuring the targeted networks in question, and though not optimal, the data used here were sufficient to address our questions of interest.

In addition, recent work by Guell and colleagues suggested that the human dentate may optimally be divided into three unique functional sub-regions (Guell et al. 2020). To this point, there are no anatomical data to support this three-way dissociation and it is based on fcMRI data; however, such a dissociation with white matter is also likely to be highly technically challenging. Because of this, we focused only on the dorsal and ventral regions as they have been mapped anatomically; however, such a parcellation in the aging brain represents an interesting future direction for work in this field.

Most notably, though we investigated sex differences in connectivity, we did not investigate menopause or have information about hormone levels in this sample. Menopause is associated with changes in the brain (e.g., Morrison et al. 2006; Robertson et al. 2009; Weber et al. 2013). Specifically, there is evidence to indicate that estrogen may have neuroprotective effects (Erickson et al. 2005; Boccardi et al. 2006; Robertson et al. 2009), and this includes the cerebellum. Further, we know that estrogen acts on the cerebellum (Hedges et al. 2012), and the hormonal changes associated with menopause may in fact be contributing to the sex differences in connectivity-age relationships seen here. Indeed, recent work looking at hormonal fluctuations during the menstrual cycle has demonstrated that over the course of the menstrual cycle, as sex steroid hormones fluctuate, increases in network coherence are related to levels of estradiol, and when progesterone rose, network coherence declines (Pritschet et al. 2020). Interestingly, when investigating the cerebellum, the dynamics were changed wherein the impacts of progesterone were larger, and those for estradiol were minimal as compared to the cortex (Fitzgerald et al. 2020). While the menstrual cycle is distinct from the hormonal changes of menopause and associated hormonal milieu, this work indicates that both cortical and cerebellar functional connectivity are sensitive to circulating sex steroid hormones. As such, f more detailed work across adulthood, taking into account reproductive status, and, optimally, sex steroid hormone levels, is warranted.

Finally, our investigation here focused on the dentate nucleus, an area of the brain with a high degree of iron (Popescu et al. 2009). The presence of iron can in turn influence the BOLD signal and, in turn, resting state connectivity measures. Furthermore, there is substantial evidence to indicate that iron concentrations differ in older adults (Rodrigue et al. 2011; Bilgic et al. 2012; Hagemeier et al. 2012; Daugherty and Raz 2015). In this sample, we were unable to quantify iron content, and we cannot rule out that this may be influencing our results. That is, iron in the dentate may actually be driving these connectivity differences, or may represent an underlying reason driving these age-connectivity patterns. The inclusion of measures of iron content in future work will be critical to better understand the impact that this may be playing on dentate connectivity, but, given that this is a key output region for the cerebellum, this will also be important for cerebellar connectivity more generally, in advanced age.

## Conclusions

Using a large sample of adults ranging from young to older adulthood, we investigated the connectivity patterns of the dorsal and ventral aspects of the human dentate nucleus. We also investigated the dissociability of these networks when using data processing methods that were designed for whole-brain analysis, as opposed to cerebellum-specific techniques. Finally, we explored sex differences in connectivity as well as relationships with age. First, using processing and analysis approaches designed for the whole-brain, we demonstrated that the dorsal and ventral dentate are associated with motor and pre-motor, and frontal and association cortices, respectively. This was further confirmed with seed contrasts, and when using a semi-partial correlation approach. With advanced age, in both the dorsal and ventral dentate regions, as age increased, connectivity with the cortex was lower, on average. Finally, we also demonstrated sex differences in dentate connectivity, as well as differences in the relationships with age when looking at males and females separately. Together, this work provides important new insights into the interactions between the cerebellar dentate nucleus and sex differences in cerebellar connectivity across the adult lifespan. This serves as a key point of comparison for studies investigating this region and associated networks in age-related disease, and for investigations incorporating needed behavioral insights.

## Acknowledgments

Data collection and sharing for this project was provided by the Cambridge Centre for Ageing and Neuroscience (CamCAN). CamCAN funding was provided by the UK Biotechnology and Biological Sciences Research Council (grant number BB/H008217/1), together with support from the UK Medical Research Council and University of Cambridge, UK. This work was further supported by R01AG065010 to J.A.B.

## References

Adams JN, Maass A, Harrison TM, Baker SL, Jagust WJ. 2019. Cortical tau deposition follows patterns of entorhinal functional connectivity in aging. Elife. 8:1–22.

Alaerts K, Swinnen SP, Wenderoth N. 2016. Sex differences in autism: A resting-state fMRI investigation of functional brain connectivity in males and females. Soc Cogn Affect Neurosci. 11:1002–1016.

Andrews-Hanna JR, Snyder AZ, Vincent JL, Lustig C, Head D, Raichle ME, Buckner RL. 2007. Disruption of large-scale brain systems in advanced aging. Neuron. 56:924–935.

Bernard JA, Goen JRM, Maldonado T. 2017. A case for motor network contributions to schizophrenia symptoms: Evidence from resting-state connectivity. Hum Brain Mapp. 38:4535–4545.

Bernard JA, Leopold DR, Calhoun VD, Mittal VA. 2015a. Regional Cerebellar Volume and Cognitive Function From Adolescence to Late Middle Age. Hum Brain Mapp. 1120:1102–1120.

Bernard JA, Leopold DR, Calhoun VD, Mittal VA. 2015b. Regional cerebellar volume and cognitive function from adolescence to late middle age. Hum Brain Mapp. 36.

Bernard JA, Orr JM, Mittal VA. 2016. Differential motor and prefrontal cerebello-cortical network development: Evidence from multimodal neuroimaging. Neuroimage. 124:591–601.

Bernard JA, Peltier SJ, Benson BL, Wiggins JL, Jaeggi SM, Buschkuehl M, Jonides J, Monk CS, Seidler RD. 2014. Dissociable functional networks of the human dentate nucleus. Cereb Cortex. 24.

Bernard JA, Peltier SJ, Wiggins JL, Jaeggi SM, Buschkuehl M, Fling BW, Kwak Y, Jonides J, Monk CS, Seidler RD. 2013. Disrupted cortico-cerebellar connectivity in older adults. Neuroimage. 83.

Bernard JA, Seidler RD. 2013. Relationships between regional cerebellar volume and sensorimotor and cognitive function in young and older adults. Cerebellum. 12.

Bernard JA, Seidler RD. 2014. Moving forward: Age effects on the cerebellum underlie cognitive and motor declines. Neurosci Biobehav Rev. 42:193–207.

Bernard JA, Seidler RD, Hassevoort KM, Benson BL, Welsh RC, Wiggins JL, Jaeggi SM, Buschkuehl M, Monk CS, Jonides J, Peltier SJ. 2012. Resting state cortico-cerebellar functional connectivity networks: a comparison of anatomical and self-organizing map approaches. Front Neuroanat. 6:1–19.

Bilgic B, Pfefferbaum A, Rohlfing T, Sullivan E V., Adalsteinsson E. 2012. MRI estimates of brain iron concentration in normal aging using quantitative susceptibility mapping. Neuroimage. 59:2625–2635.

Biswal BB, Mennes M, Zuo X-N, Gohel S, Kelly C, Smith SM, Beckmann CF, Adelstein JS, Buckner RL, Colcombe S, Dogonowski A-M, Ernst M, Fair D, Hampson M, Hoptman MJ, Hyde JS, Kiviniemi VJ, Kötter R, Li S-J, Lin C-P, Lowe MJ, Mackay C, Madden DJ, Madsen KH, Margulies DS, Mayberg HS, McMahon K, Monk CS, Mostofsky SH, Nagel BJ, Pekar JJ, Peltier SJ, Petersen SE, Riedl V, Rombouts S a RB, Rypma B, Schlaggar BL, Schmidt S, Seidler RD, Siegle GJ, Sorg C, Teng G-J, Veijola J, Villringer A, Walter M, Wang L, Weng X-C, Whitfield-Gabrieli S, Williamson P, Windischberger C, Zang Y-F, Zhang H-Y, Castellanos FX, Milham MP. 2010. Toward discovery science of human brain function. Proc Natl Acad Sci U S A. 107:4734–4739.

Bo J, Lee C-M, Kwak Y, Peltier SJ, Bernard JA, Buschkuehl M, Jaeggi SM, Wiggins JL, Jonides J, Monk CS, Seidler RD. 2014. Lifespan differences in cortico-striatal resting state connectivity. Brain Connect. 4.

Boccardi M, Ghidoni R, Govoni S, Testa C, Benussi L, Bonetti M, Binetti G, Frisoni GB. 2006. Effects of hormone therapy on brain morphology of healthy postmenopausal women: A Voxel-based morphometry study. Menopause. 13:584–591.

Buckner RL, Krienen FM, Castellanos A, Diaz JC, Thomas Yeo BT. 2011. The organization of the human cerebellum estimated by intrinsic functional connectivity. J Neurophysiol. 106:2322–2345.

Cabeza R. 2002. Hemispheric asymmetry reduction in older adults: The HAROLD model. Psychol Aging. 17:85–100.

Cabeza R, Albert M, Belleville S, Craik FIM, Duarte A, Grady CL, Lindenberger U, Nyberg L, Park DC, Reuter-lorenz PA, Rugg MD, Steffener J. 2018. neuroscience of healthy ageing. Nat Rev Neurosci.

Cabeza R, Anderson ND, Locantore JK, McIntosh AR. 2002. Aging Gracefully: Compensatory Brain Activity in High-Performing Older Adults. Neuroimage. 17:1394–1402.

Carter CL, Resnick EM, Mallampalli M, Kalbarczyk A. 2012. Sex and gender differences in Alzheimer’s disease: Recommendations for future research. J Women’s Heal. 21:1018–1023.

Daugherty AM, Raz N. 2015. Appraising the Role of Iron in Brain Aging and Cognition: Promises and Limitations of MRI Methods. Neuropsychol Rev. 25:272–287.

Diedrichsen J. 2006. A spatially unbiased atlas template of the human cerebellum. Neuroimage. 33:127–138.

Diedrichsen J, Balsters JH, Flavell J, Cussans E, Ramnani N. 2009. A probabilistic MR atlas of the human cerebellum. Neuroimage. 46:39–46.

Dum RP, Strick PL. 2003a. An Unfolded Map of the Cerebellar Dentate Nucleus and its Projections to the Cerebral Cortex. J Neurophysiol. 89:634–639.

Dum RP, Strick PL. 2003b. An unfolded map of the cerebellar dentate nucleus and its projections to the cerebral cortex. J Neurophysiol. 89:634–639.

Engman J, Linnman C, Van Dijk KRA, Milad MR. 2016. Amygdala subnuclei resting-state functional connectivity sex and estrogen differences. Psychoneuroendocrinology. 63:34–42.

Erickson KI, Colcombe SJ, Raz N, Korol DL, Scalf P, Webb A, Cohen NJ, McAuley E, Kramer AF. 2005. Selective sparing of brain tissue in postmenopausal women receiving hormone replacement therapy. Neurobiol Aging. 26:1205–1213.

Fearnley JM, Lees AJ. 1991. Ageing and Parkinson’S Disease◻: Substantia Nigra Regional Selectivity. Brain, A J Neurol. 114:2283–2301.

Ferreira LK, Busatto GF. 2013. Resting-state functional connectivity in normal brain aging. Neurosci Biobehav Rev. 37:384–400.

Ferreira LK, Regina ACB, Kovacevic N, Martin MDGM, Santos PP, Carneiro CDG, Kerr DS, Amaro E, Mcintosh AR, Busatto GF. 2016. Aging effects on whole-brain functional connectivity in adults free of cognitive and psychiatric disorders. Cereb Cortex. 26:3851–3865.

Fitzgerald M, Pritschet L, Santander T, Grafton ST, Jacobs EG. 2020. Cerebellar network organization across the human menstrual cycle. Sci Rep. 10:1–11.

Gellersen HM, Guell X, Sami S. 2020. Differential vulnerability of the cerebellum in healthy ageing and Alzheimer’s disease. medRxiv.

Guell X, D’Mello AM, Hubbard NA, Romeo RR, Gabrieli JDE, Whitfield-Gabrieli S, Schmahmann JD, Anteraper SA. 2020. Functional Territories of Human Dentate Nucleus. Cereb Cortex. 30:2401–2417.

Guell X, Schmahmann JD, Gabrieli JDE, Ghosh SS. 2018. Functional gradients of the cerebellum. Elife. 7:1–22.

Guo CC, Tan R, Hodges JR, Hu X, Sami S. 2016. Network-selective vulnerability of the human cerebellum to Alzheimer’s disease and frontotemporal dementia. 1527–1538.

Hagemeier J, Geurts JJG, Zivadinov R. 2012. Brain iron accumulation in aging and neurodegenerative disorders. Expert Rev Neurother. 12:1467–1480.

Han S, An Y, Carass A, Prince JL, Resnick SM. 2020. NeuroImage Longitudinal analysis of regional cerebellum volumes during normal aging. Neuroimage. 220:117062.

Hartholt KA, Van Beeck EF, Polinder S, Van Der Velde N, Van Lieshout EMM, Panneman MJM, Van Der Cammen TJM, Patka P. 2011. Societal consequences of falls in the older population: Injuries, healthcare costs, and long-term reduced quality of life. J Trauma – Inj Infect Crit Care. 71:748–753.

Hausman HK, Jackson TB, Goen JRM, Bernard JA. 2020. From Synchrony to Asynchrony: Cerebellar-Basal Ganglia Functional Circuits in Young and Older Adults. Cereb Cortex. 30:718–729.

Hedges VL, Ebner TJ, Meisel RL, Mermelstein PG. 2012. The cerebellum as a target for estrogen action. Front Neuroendocrinol. 33:403–411.

Jacobs HIL, Hopkins DA, Mayrhofer HC, Bruner E, Leeuwen FW Van, Raaijmakers W, Schmahmann JD. 2018. The cerebellum in Alzheimer’s disease◻: evaluating its role in cognitive decline. 37–47.

Kelly C, De Zubicaray G, Di Martino A, Copland DA, Reiss PT, Klein DF, Castellanos FX, Milham MP, McMahon K. 2009. L-dopa modulates functional connectivity in striatal cognitive and motor networks: A double-blind placebo-controlled study. J Neurosci. 29:7364–7378.

Kelly RM, Strick PL. 2003. Cerebellar Loops with Motor Cortex and Prefrontal Cortex of a Nonhuman Primate. J Neurosci. 23:8432–8444.

King M, Hernandez-castillo C, Sereno M, Ivry RB. 2019. Review Universal Transform or Multiple Functionality◻? Understanding the Contribution of the Human Cerebellum across Task Domains.

King M, Hernandez-castillo CR, Poldrack RA, Ivry RB, Diedrichsen J. 2019. Functional boundaries in the human cerebellum revealed by a multi-domain task battery. Nat Neurosci. 22:1371–1378.

Koppelmans V, Hirsiger S, Susan M, Seidler RD. 2015. Cerebellar Gray and White Matter Volume and Their Relation With Age and Manual Motor Performance in Healthy Older Adults. 2363:2352–2363.

Koppelmans V, Hoogendam YY, Hirsiger S, Mérillat S, Jäncke L, Seidler RD. 2017. Regional cerebellar volumetric correlates of manual motor and cognitive function. Brain Struct Funct. 222:1929–1944.

Krienen FM, Buckner RL. 2009. Segregated Fronto-Cerebellar Circuits Revealed by Intrinsic Functional Connectivity. Cereb Cortex. 19:2485–2497.

Lin CY, Chen CH, Tom SE, Kuo SH. 2020. Cerebellar Volume Is Associated with Cognitive Decline in Mild Cognitive Impairment: Results from ADNI. Cerebellum. 19:217–225.

MacLullich AMJ, Edmond CL, Ferguson KJ, Wardlaw JM, Starr JM, Seckl JR, Deary IJ. 2004. Size of the neocerebellar vermis is associated with cognition in healthy elderly men. Brain Cogn. 56:344–348.

Marek S, Siegel JS, Gordon EM, Raut R V., Gratton C, Newbold DJ, Ortega M, Laumann TO, Adeyemo B, Miller DB, Zheng A, Lopez KC, Berg JJ, Coalson RS, Nguyen AL, Dierker D, Van AN, Hoyt CR, McDermott KB, Norris SA, Shimony JS, Snyder AZ, Nelson SM, Barch DM, Schlaggar BL, Raichle ME, Petersen SE, Greene DJ, Dosenbach NUF. 2018. Spatial and Temporal Organization of the Individual Human Cerebellum. Neuron. 100:977–993.e7.

Mazure CM, Swendsen J. 2016. Sex differences in Alzheimer’s disease and other dementias. Lancet Neurol. 15:451–452.

McGeer PL, McGeer EG, Suzuki JS. 1977. Aging and Extrapyramidal Function. Arch Neurol. 34:33–35.

Miller TD, Ferguson KJ, Reid LM, Wardlaw JM, Starr JM, Seckl JR, Deary IJ, Maclullich AMJ. 2013. Cerebellar vermis size and cognitive ability in community-dwelling elderly men. Cerebellum. 12:68–73.

Morrison JH, Brinton RD, Schmidt PJ, Gore AC. 2006. Estrogen, Menopause, and the Aging Brain: How Basic Neuroscience Can Inform Hormone Therapy in Women. J Neurosci. 26:10332–10348.

O’Reilly JX, Beckmann CF, Tomassini V, Ramnani N, Johansen-Berg H. 2010. Distinct and overlapping functional zones in the cerebellum defined by resting state functional connectivity. Cereb Cortex. 20:953–965.

Oguro H, Okada K, Yamaguchi S, Kobayashi S. 1998. Sex differences in morphology of the brain stem and cerebellum with normal ageing. Neuroradiology. 40:788–792.

Olivito G, Serra L, Marra C, Di Domenico C, Caltagirone C, Toniolo S, Cercignani M, Leggio M, Bozzali M. 2020. Cerebellar dentate nucleus functional connectivity with cerebral cortex in Alzheimer’s disease and memory: a seed-based approach. Neurobiol Aging. 89:32–40.

Oostland M, van Hooft J. 2016. Serotonin in the cerebellum. In: Essentials of Cerebellum and Cerebellar Disorders. p. 243–247.

Oschmann M, Gawryluk JR. 2020. A Longitudinal Study of Changes in Resting-State Functional Magnetic Resonance Imaging Functional Connectivity Networks during Healthy Aging. Brain Connect. 10:377–384.

Park DC, Polk TA, Mikels JA, Taylor SF, Marshuetz C. 2001. Cerebral aging: integration of brain and behavioral models of cognitive function. Dialogues Clin Neurosci. 3:151–165.

Park DC, Reuter-Lorenz P. 2009. The adaptive brain: aging and neurocognitive scaffolding. Annu Rev Psychol. 60:173–196.

Popescu BFG, Robinson CA, Rajput A, Rajput AH, Harder SL, Nichol H. 2009. Iron, copper, and zinc distribution of the cerebellum. Cerebellum. 8:74–79.

Power JD, Barnes K a, Snyder AZ, Schlaggar BL, Petersen SE. 2012. Spurious but systematic correlations in functional connectivity MRI networks arise from subject motion. Neuroimage. 59:2142–2154.

Pritschet L, Santander T, Taylor CM, Layher E, Yu S, Miller MB, Grafton ST, Jacobs EG. 2020. Functional reorganization of brain networks across the human menstrual cycle. Neuroimage. 220:117091.

Puts MTE, Lips P, Deeg DJH. 2005. Sex differences in the risk of frailty for mortality independent of disability and chronic diseases. J Am Geriatr Soc. 53:40–47.

Ramnani N. 2006. The primate cortico-cerebellar system: anatomy and function. Nat Rev Neurosci. 7:511–522.

Raz N, Dupuis JH, Briggs SD, McGavran C, Acker JD. 1998. Differential effects of age and sex on the cerebellar hemispheres and the vermis: a prospective MR study. AJNR Am J Neuroradiol. 19:65–71.

Raz N, Ghisletta P, Rodrigue KM, Kennedy KM, Lindenberger U. 2010. Trajectories of brain aging in middle-aged and older adults: regional and individual differences. Neuroimage. 51:501–511.

Raz N, Gunning-Dixon F, Head D, Williamson a, Acker JD. 2001. Age and sex differences in the cerebellum and the ventral pons: a prospective MR study of healthy adults. AJNR Am J Neuroradiol. 22:1161–1167.

Raz N, Gunning FM, Head D, Dupuis JH, McQuain J, Briggs SD, Loken WJ, Thornton a E, Acker JD. 1997. Selective aging of the human cerebral cortex observed in vivo: differential vulnerability of the prefrontal gray matter. Cereb Cortex. 7:268–282.

Raz N, Lindenberger U, Rodrigue KM, Kennedy KM, Head D, Williamson A, Dahle C, Gerstorf D, Acker JD. 2005. Regional brain changes in aging healthy adults: general trends, individual differences and modifiers. Cereb Cortex. 15:1676–1689.

Raz N, Schmiedek F, Rodrigue KM, Kennedy KM, Lindenberger U, Lövdén M. 2013. Differential brain shrinkage over 6months shows limited association with cognitive practice. Brain Cogn. 82:171–180.

Reuter-Lorenz P a., Cappell K a. 2008. Neurocognitive Aging and the Compensation Hypothesis. Curr Dir Psychol Sci. 17:177–182.

Reuter-Lorenz P a., Stanczak L, Miller a. C. 1999. Neural Recruitment and Cognitive Aging: Two Hemispheres Are Better Than One, Especially as You Age. Psychol Sci. 10:494–500.

Reuter-Lorenz PA, Park DC. 2014. How Does it STAC Up? Revisiting the Scaffolding Theory of Aging and Cognition. Neuropsychol Rev. 24:355–370.

Robertson D, Craig M, Van Amelsvoort T, Daly E, Moore C, Simmons A, Whitehead M, Morris R, Murphy D. 2009. Effects of estrogen therapy on age-related differences in gray matter concentration. Climacteric. 12:301–309.

Rodrigue KM, Haacke EM, Raz N. 2011. Differential effects of age and history of hypertension on regional brain volumes and iron. Neuroimage. 54:750–759.

Saccà V, Sarica A, Quattrone A, Rocca F, Quattrone A, Novellino F. 2021. Aging effect on head motion: A Machine Learning study on resting state fMRI data. J Neurosci Methods. 352.

Salmi J, Pallesen K, Neuvonen T. 2010. Cognitive and motor loops of the human cerebro-cerebellar system. J Cogn …. 2663–2676.

Scheinost D, Finn ES, Tokoglu F, Shen X, Papademetris X, Hampson M, Constable RT. 2015. Sex differences in normal age trajectories of functional brain networks. Hum Brain Mapp. 36:1524–1535.

Schmahmann JD. 2018. The cerebellum and cognition. Neurosci Lett. 62–75.

Schmahmann JD, Sherman JC. 1998. The cerebellar cognitive affective syndrome. Brain. 121:561–579.

Seidler RD, Bernard JA, Burutolu TB, Fling BW, Gordon MT, Gwin JT, Kwak Y, Lipps DB. 2010a. Motor control and aging: Links to age-related brain structural, functional, and biochemical effects. Neurosci Biobehav Rev. 34.

Seidler RD, Bernard JA, Burutolu TB, Fling BW, Gordon MT, Gwin JT, Kwak Y, Lipps DB. 2010b. Motor control and aging: links to age-related brain structural, functional, and biochemical effects. Neurosci Biobehav Rev. 34:721–733.

Sereno MI, Diedrichsen J, Tachrount M, Testa-silva G. 2020. The human cerebellum has almost 80 % of the surface area of the neocortex. 1–6.

Shafto MA, Tyler LK, Dixon M, Taylor JR, Rowe JB, Cusack R, Calder AJ, Marslen-Wilson WD, Duncan J, Dalgleish T, Henson RN, Brayne C, Bullmore E, Campbell K, Cheung T, Davis S, Geerligs L, Kievit R, McCarrey A, Price D, Samu D, Treder M, Tsvetanov K, Williams N, Bates L, Emery T, Erzinçlioglu S, Gadie A, Gerbase S, Georgieva S, Hanley C, Parkin B, Troy D, Allen J, Amery G, Amunts L, Barcroft A, Castle A, Dias C, Dowrick J, Fair M, Fisher H, Goulding A, Grewal A, Hale G, Hilton A, Johnson F, Johnston P, Kavanagh-Williamson T, Kwasniewska M, McMinn A, Norman K, Penrose J, Roby F, Rowland D, Sargeant J, Squire M, Stevens B, Stoddart A, Stone C, Thompson T, Yazlik O, Barnes D, Hillman J, Mitchell J, Villis L, Matthews FE. 2014. The Cambridge Centre for Ageing and Neuroscience (Cam-CAN) study protocol: A cross-sectional, lifespan, multidisciplinary examination of healthy cognitive ageing. BMC Neurol. 14:1–25.

Smith REW, Avery JA, Wallace GL, Kenworthy L, Gotts SJ, Martin A. 2019. Sex differences in resting-state functional connectivity of the cerebellum in autism spectrum disorder. Front Hum Neurosci. 13:1–13.

Steele CJ, Anwander A, Bazin P, Trampel R, Schaefer A, Turner R, Ramnani N, Villringer A. 2017. Human Cerebellar Sub-millimeter Diffusion Imaging Reveals the Motor and Non-motor Topography of the Dentate Nucleus. 4537–4548.

Steele CJ, Chakravarty MM. 2017. Gray-matter structural variability in the human cerebellum: Lobule-specific differences across sex and hemisphere. Neuroimage.

Steele CJ, Chakravarty MM. 2018. Gray-matter structural variability in the human cerebellum: Lobule-specific differences across sex and hemisphere. Neuroimage. 170:164–173.

Stevens J a, Sogolow ED. 2005. Gender differences for non-fatal unintentional fall related injuries among older adults. Inj Prev. 11:115–119.

Stoodley CJ, Valera EM, Schmahmann JD. 2012. Functional topography of the cerebellum for motor and cognitive tasks: An fMRI study. Neuroimage. 59:1560–1570.

Strick PL, Dum RP, Fiez JA. 2009. Cerebellum and Nonmotor Function. Annu Rev Neurosci. 32:413–434.

Tabatabaei-Jafari H, Walsh E, Shaw ME, Cherbuin N. 2017. The cerebellum shrinks faster than normal ageing in Alzheimer’s disease but not in mild cognitive impairment. Hum Brain Mapp. 00.

Taylor JR, Williams N, Cusack R, Auer T, Shafto MA, Dixon M, Tyler LK, Cam-CAN, Henson RN. 2017. The Cambridge Centre for Ageing and Neuroscience (Cam-CAN) data repository: Structural and functional MRI, MEG, and cognitive data from a cross-sectional adult lifespan sample. Neuroimage. 144:262–269.

Tiemeier H, Lenroot RK, Greenstein DK, Tran L, Pierson R, Giedd JN. 2010. Cerebellum development during childhood and adolescence: A longitudinal morphometric MRI study. Neuroimage. 49:63–70.

Tomasi D, Volkow ND. 2012. Aging and functional brain networks. Mol Psychiatry. 17:549–558.

Toniolo S, Serra L, Olivito G, Marra C, Bozzali M. 2018. Patterns of Cerebellar Gray Matter Atrophy Across Alzheimer’s Disease Progression. 12:1–8.

Van Dijk KR a, Sabuncu MR, Buckner RL. 2012. The influence of head motion on intrinsic functional connectivity MRI. Neuroimage. 59:431–438.

Weber MT, Rubin LH, Maki PM. 2013. Cognition in perimenopause: the effect of transition stage. Menopause. 20:511–517.

Weis S, Hodgetts S, Hausmann M. 2019. Sex differences and menstrual cycle effects in cognitive and sensory resting state networks. Brain Cogn. 131:66–73.

Whitfield-Gabrieli S, Nieto Castañón A. 2012. Conn: a functional connectivity toolbox for correlated and anticorrelated brain networks. Brain Connect. 2:125–141.

Zheng W, Liu X, Song H, Li K, Wang Z. 2017. Altered functional connectivity of cognitive-related cerebellar subregions in alzheimer’s disease. Front Aging Neurosci. 9:1–9.

