## Supplemental Tables for "Cerebellar Dentate Connectivity Across Adulthood: A Large-Scale Resting State Functional Connectivity Investigation"

**Supplementary Table 1.** Average (and standard deviation) of mean motion and maximum motion by age decade, with the sample size indicated in parentheses.

| **Age Group** | **Mean Motion (mm)** | **Maximum Motion (mm)** |
| --- | --- | --- |
| 18-29 (n=67) | .173 (.050) | .938 (1.58) |
| 30-39 (n=87) | .181 (.061) | .734 (.635) |
| 40-49 (n=96) | .211 (.072) | .772 (.483) |
| 50-59 (n=87) | .224 (.068) | 1.11 (.655) |
| 60-69 (n=93) | .248 (.072) | 1.06 (1.04) |
| 70-79 (n=100) | .270 (.067) | 1.42 (1.07) |
| 80-89 (n=60) | .271 (.068) | 1.52 (.903) |

**Supplementary Table 2.** Dorsal and Ventral Dentate Connectivity. Anatomical locations determined using Harvard-Oxford max probability atlas.

| **Region** | **Cluster Size** | **MNI Coordinates** | | | **P(FDR)** | **Beta** |
| --- | --- | --- | --- | --- | --- | --- |
|  |  | **X** | **Y** | **Z** |  |  |
| **DORSAL SEED** | | | | | | |
| **CB** | 51989 | 12 | -58 | -30 | 0.0000 | 0.0500 |
| Temporal fusiform cortex, posterior | 43579 | -36 | -34 | -6 | 0.0000 | -0.0280 |
| Middle frontal gyrus | 1699 | -42 | 0 | 62 | 0.0000 | 0.0230 |
| Anterior cingulate gyrus | 787 | 4 | 28 | 24 | 0.0000 | 0.0230 |
| Middle frontal gyrus | 569 | 36 | 8 | 66 | 0.0000 | 0.0230 |
| Frontal orbital cortex | 177 | -24 | 32 | -16 | 0.0001 | 0.0210 |
| Superior frontal gyrus | 88 | 0 | -48 | 32 | 0.0046 | -0.0180 |
| Frontal pole | 69 | 36 | 42 | 44 | 0.0127 | 0.0230 |
| **VENTRAL SEED** | | | | | | |
| Supplementary motor cortex (juxtapositional cortex) | 70020 | 2 | -4 | 48 | 0.0000 | -0.0340 |
| Lobule VI/Crus I | 56726 | 18 | -66 | -36 | 0.0000 | 0.0560 |
| Middle temporal gyrus | 6109 | 60 | -38 | -2 | 0.0000 | 0.0280 |
| Posterior cingulate gyrus | 205 | -20 | -54 | -64 | 0.0000 | -0.0230 |
| Caudate | 156 | 18 | 8 | 14 | 0.0001 | 0.0230 |
| Lateral occipital cortex | 79 | -44 | -86 | 26 | 0.0046 | -0.0220 |
| Frontal pole | 74 | 20 | 62 | 0 | 0.0054 | -0.0220 |
| Brainstem/white matter | 65 | -6 | -24 | -58 | 0.0085 | -0.0200 |
| Amygdala | 41 | 30 | -4 | -20 | 0.0423 | 0.0200 |
| **VENTRAL > DORSAL** | | | | | | |
| Dentate | 44678 | 18 | -66 | -36 | 0.0000 | 0.0450 |
| Middle temporal gyrus | 1631 | 60 | -32 | -4 | 0.0000 | 0.0290 |
| Inferior frontal gyrus, *pars triangularis* | 780 | 56 | 34 | 16 | 0.0000 | 0.0280 |
| Temporal pole | 306 | 60 | 8 | -22 | 0.0000 | 0.0290 |
| Thalamus | 57 | -8 | -28 | 6 | 0.0278 | 0.0250 |
| Lingual gyrus | 49 | 12 | -64 | -2 | 0.0389 | 0.0240 |
| **DORSAL > VENTRAL** | | | | | | |
| Dentate/interposed nuclei | 60181 | 12 | -58 | -30 | 0.0000 | 0.0410 |
| Frontal pole | 1608 | 34 | 46 | 32 | 0.0000 | 0.0330 |
| Middle frontal gyrus | 481 | -28 | 36 | 28 | 0.0000 | 0.0300 |
| Occipital pole | 431 | -22 | -98 | -20 | 0.0000 | 0.0280 |
| Occipital pole | 98 | 22 | -100 | -12 | 0.0020 | 0.0260 |
| Frontal pole | 65 | 24 | 62 | -8 | 0.0105 | 0.0260 |

**Supplementary Table 3.** Patterns of connectivity for the dorsal and ventral dentate when analyzed using a semi-partial correlation approach, wherein we controlled for the signal in the other seed.

| **Region** | **Cluster Size** | **MNI Coordinates** | | | **P(FDR)** | **Beta** |
| --- | --- | --- | --- | --- | --- | --- |
|  |  | **X** | **Y** | **Z** |  |  |
| **DORSAL SEED** | | | | | | |
| Lingual gyrus | 51658 | 12 | -58 | -30 | 0.0000 | 0.0490 |
| Temporal fusiform cortex | 37878 | -36 | -34 | -6 | 0.0000 | -0.0290 |
| Anterior cingulate gyrus | 1489 | 2 | 28 | 28 | 0.0000 | 0.0270 |
| Precentral gyrus | 1032 | -50 | -6 | 56 | 0.0000 | 0.0250 |
| Middle frontal gyrus (dorsal pre-motor) | 1029 | 36 | 8 | 66 | 0.0000 | 0.0240 |
| Frontal orbital cortex | 113 | -24 | 32 | -16 | 0.0009 | 0.0240 |
| Frontal pole | 56 | 24 | 62 | -10 | 0.0273 | 0.0230 |
| **VENTRAL SEED** | | | | | | |
| Juxtapositional cortex (supplementary motor cortex) | 68223 | 2 | 6 | 46 | 0.0000 | -0.0360 |
| Dentate nucleus | 53624 | 18 | -66 | -36 | 0.0000 | 0.0570 |
| Middle temporal gyrus | 5678 | 60 | -38 | -2 | 0.0000 | 0.0300 |
| Lobule VIIIb | 570 | -20 | -56 | -64 | 0.0000 | -0.0270 |
| Lobule VIIIa | 161 | 32 | -52 | -62 | 0.0001 | -0.0240 |
| Brainstem | 120 | 22 | -16 | -6 | 0.0004 | 0.0220 |
| Caudate | 104 | 18 | 8 | 14 | 0.0008 | 0.0230 |
| Frontal pole | 76 | 20 | 62 | 0 | 0.0038 | -0.0240 |
| Brainstem | 62 | -6 | -24 | -58 | 0.0084 | -0.0210 |
| Midbrain | 54 | -6 | -8 | -10 | 0.0130 | -0.0240 |
| Temporal pole | 53 | 28 | 8 | -50 | 0.0130 | -0.0230 |
| Caudate | 45 | -18 | 26 | 0 | 0.0216 | -0.0220 |
| Lateral occipital cortex | 43 | -44 | -86 | 26 | 0.0233 | -0.0240 |
| Hypothalamus | 38 | 4 | -2 | -4 | 0.0321 | 0.0210 |
| Amygdala | 33 | 30 | -4 | -20 | 0.0453 | 0.0220 |

**Supplementary Table 4.** Quadratic relationships between dentate connectivity and age. With the positive relationships, this can be imagined as analogous to an “inverted-U” pattern, while the negative follow that of a “u”.

| **Region** | **Cluster Size** | **MNI Coordinates** | | | **P(FDR)** |
| --- | --- | --- | --- | --- | --- |
|  |  | **X** | **Y** | **Z** |  |
| **DORSAL SEED** | | | | | |
| **Positive Relationships** | | | | | |
| Lobule V | 120 | 18 | -48 | -20 | 0.007 |
| Vermis VI | 117 | -4 | -64 | -28 | 0.007 |
| Lobule V | 75 | 22 | -46 | -30 | 0.037 |
| **Negative Relationships** | | | | | |
| Paracingulate gyrus | 160 | 6 | 50 | 10 | 0.003 |
| Precuneous | 108 | 0 | -64 | 34 | 0.015 |
| Frontal orbital cortex | 77 | 50 | 22 | -8 | 0.048 |
| **VENTRAL SEED** | | | | | |
| N/A | | | | | |

**Supplementary Table 5.** Quadratic relationships between dentate connectivity and age in females. With the positive relationships, this can be imagined as analogous to an “inverted-U” pattern, while the negative follow that of a “u”.

| **Region** | **Cluster Size** | **MNI Coordinates** | | | **P(FDR)** |
| --- | --- | --- | --- | --- | --- |
|  |  | **X** | **Y** | **Z** |  |
| **DORSAL SEED** | | | | | |
| **Positive** | | | | | |
| N/A | | | | | |
| **Negative** | | | | | |
| Anterior cingulate gyrus | 150 | 12 | 40 | 16 | 0.003 |
| **VENTRAL SEED** | | | | | |
| N/A | | | | | |

**Supplementary Table 6.** Quadratic relationships between dentate connectivity and age in males. With the positive relationships, this can be imagined as analogous to an “inverted-U” pattern, while the negative follow that of a “u”.

| **Region** | **Cluster Size** | **MNI Coordinates** | | | **P(FDR)** |
| --- | --- | --- | --- | --- | --- |
|  |  | **X** | **Y** | **Z** |  |
| **DORSAL SEED** | | | | | |
| **Positive** | | | | | |
| Lobule VI | 1074 | 24 | -68 | -26 | 0 |
| Lobule VI | 140 | -8 | -68 | -24 | 0.001 |
| Lobule VI | 99 | -22 | -58 | -26 | 0.008 |
| Dentate/interposed nuclei | 62 | -14 | -44 | -30 | 0.045 |
| **Negative** | | | | | |
| N/A | | | | | |
| **VENTRAL SEED** | | | | | |
| **Positive** | | | | | |
| Lobule VI | 404 | 26 | -52 | -34 | 0 |
| Putamen/internal capsule | 95 | -24 | -6 | 14 | 0.018 |
| **Negative** | | | | | |
| N/A | | | | | |

**Supplementary Table 7.** Differential relationships between connectivity and age for females and males (interactions).

| **Region** | **Cluster Size** | **MNI Coordinates** | | | **P(FDR)** |
| --- | --- | --- | --- | --- | --- |
|  |  | **X** | **Y** | **Z** |  |
| **DORSAL SEED** | | | | | |
| **Females>Males** | | | | | |
| Parietal operculum cortex | 114 | 30 | -30 | 24 | 0.014 |
| **Males>Females** | | | | | |
| Lateral occipital cortex (angular gyrus) | 107 | 50 | -62 | 26 | 0.017 |
| **VENTRAL SEED** | | | | | |
| **Females>Males** | | | | | |
| Superior temporal gyrus, posterior | 109 | -58 | -30 | 2 | 0.013 |
| **Males>Females** | | | | | |
| N/A | | | | | |

**Supplementary Figure 1**. Functional connectivity patterns for the dorsal (yellow/orange) and ventral (blue/purple/pink) dentate seeds calculated when using a semi-partial correlation. The patterns of connectivity for the two seeds parallel the dissociation seen in Figure 2 when using a standard bivariate correlation approach, wherein the dorsal seed is more strongly associated with motor cortical regions, while the ventral seed is more strongly associated with frontal and association regions.

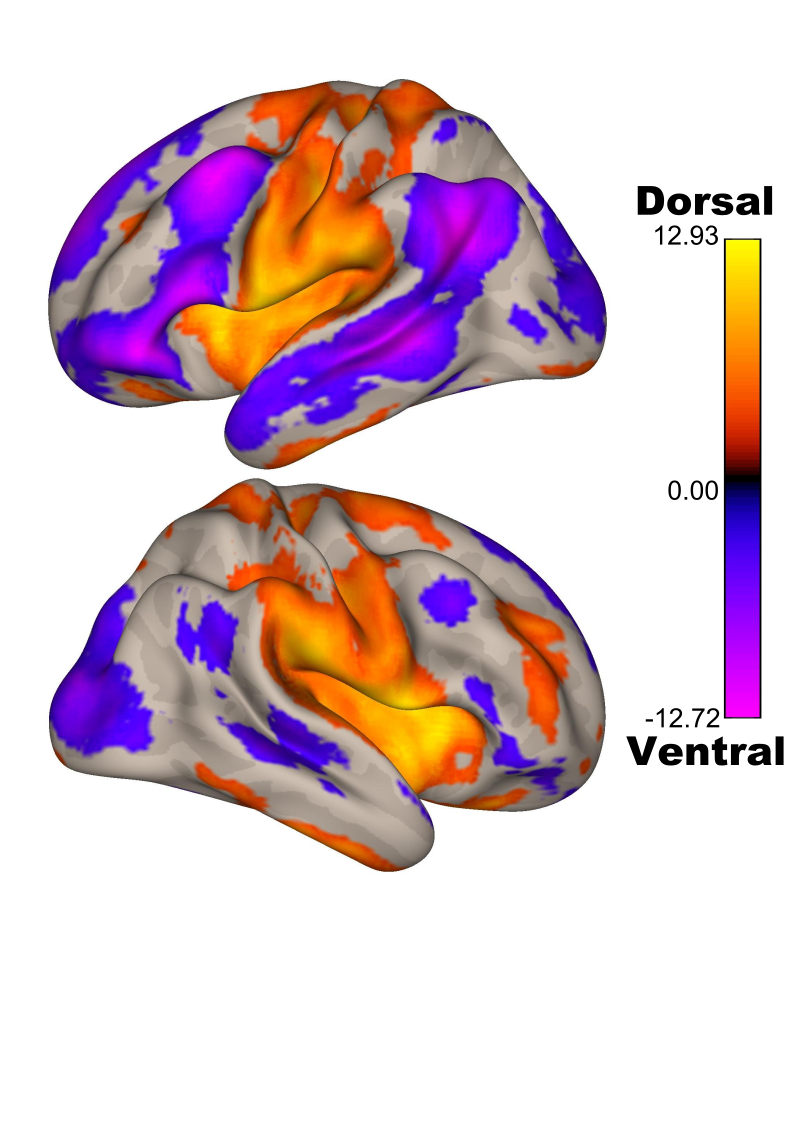
